# ‘RMT-Finder’: an automated procedure to determine the Resting Motor Threshold for Transcranial Magnetic Stimulation

**DOI:** 10.64898/2026.03.25.714235

**Authors:** Laurine F. Boidequin, Marcos Moreno-Verdú, Baptiste M. Waltzing, Julien J. Lambert, Elise E. Van Caenegem, Charlène Truong, Robert M. Hardwick

## Abstract

**Background:** Transcranial Magnetic Stimulation (TMS) studies identify the Resting Motor Threshold (RMT) to calibrate stimulation intensity. However, this procedure is time-consuming and subject to variability. We developed an automated procedure to improve the efficiency and standardization of RMT determination.

**New method:** We developed an algorithm that measures MEP amplitudes and automatically adjusts stimulation intensity to determine the RMT. Experiment 1 compared this automated method with the manual procedure in terms of reliability and equivalence. Experiment 2 developed a “Fast” automated process, assessing it against both the manual and initial automated procedures.

**Results:** Across both experiments the automated approach demonstrated excellent test-retest reliability and strong agreement with the manual method (Intraclass Correlation Coefficients ≥0.95), giving estimates of RMT statistically equivalent to those of manual measurements within ±3% MSO, with the majority of comparisons within ±2% MSO. Experiment 2 optimized the procedure, allowing empirical determination of the RMT in an average of <3 minutes with only 33-34 pulses.

**Comparison with existing methods:** ‘RMT-Finder’ provides a reliable and time-efficient alternative to manual approaches. To the best of our knowledge RMT-Finder presents the first ‘closed-loop feedback’ approach to identify the RMT without manual intervention. This procedure can improve standardization and reproducibility in TMS studies.

**Conclusions:** Automating RMT assessment allows rapid and highly reproducible assessment of this standard TMS measurement, making it viable for inclusion in routine clinical applications that require standardized procedures.

## Introduction

Transcranial Magnetic Stimulation (TMS) is a non-invasive brain stimulation technique widely used to investigate motor and cognitive processes (Hallett, 2007). Applying TMS over the primary motor cortex can produce focal muscle twitches; the magnitude of this response can be recorded using electromyography (EMG) as a Motor Evoked Potential (MEP). Given that this response can be recorded and quantified precisely, the MEP has become the standard approach to determining dose-response effects when using TMS. The intensity of TMS that is applied is typically calculated in relation to the “Resting Motor Threshold” (RMT), defined as the lowest intensity of the Maximal Stimulator Output (MSO) required to produce MEPs with a peak-to-peak amplitude of ≥50μV in at least 5 out of 10 trials (Rossini et al., 2015).

Determining the RMT using ‘relative-frequency’ methods (i.e. delivering pulses at a given intensity until it can be confirmed/rejected as a possible value for the RMT) is common practice in TMS studies (Hamoline et al., 2024; Rossini et al., 2015, 1994; Vassiliadis et al., 2020). Unfortunately this is relatively time-consuming, usually involving delivering ∼50-75 TMS pulses (Mishory et al., 2004). This has led researchers to search for more efficient approaches to determining the RMT. “Threshold tracking” methods provide a conceptually elegant alternative; the response to TMS is modelled as a cumulative gaussian distribution, allowing the RMT to be estimated using relatively few pulses delivered across a range of intensities (Awiszus, 2003). This approach can estimate the RMT in as few as 19±6 pulses (Awiszus, 2003), but it has also been reported that using relatively few measurements can lead to a substantial misestimation of the RMT (Koponen and Peterchev, 2022). Moreover, given the high intrinsic variability of MEPs (Rossini et al., 1994), relying on a small number of responses per intensity may be problematic, as threshold estimation may be disproportionately influenced by a few unusually large or small MEPs. Most researchers continue to use relative-frequency based approaches (Rossini et al., 2015), presumably as it provides direct empirical data to demonstrate the RMT (i.e. it *shows* that the intensity at −1% of the RMT fails to give the required criterion). A balance therefore needs to be found between efficiency, empirical robustness, and practicality when determining the RMT. Developing methods that optimize the speed of the process while preserving accuracy and reliability would thus represent a valuable advance.

The goal of the present study was therefore to optimize the determination of the RMT based on a “relative-frequency” approach. Given that these methods typically require a human experimenter to monitor MEP amplitudes and make decisions on how to proceed (Awiszus and Borckardt, Jeffrey J., 2011; Julkunen, 2019), we reasoned that automating this process would not only improve the speed of decision making and decrease the possibility for human error but also provide a standardized procedure that is readily transferable across laboratories. We therefore developed software that measures peak-to-peak MEP amplitudes on a trial-by-trial basis, identifies whether a given intensity of stimulation could represent the RMT, and modifies the intensity of the stimulator accordingly. This in turn allows the experimenter to focus on precise coil positioning, reducing the likelihood of human errors.We predicted that our automated procedure would provide measurements of the RMT with accuracy comparable to traditional manual estimation, but with considerably higher efficiency.

## Methods

### Participants

A total of 48 healthy participants completed the study, 24 in Experiment 1 and 24 in Experiment 2. None of them took part in both experiments (details of each sample are provided in the relevant experimental section). The experiments were conducted at the Institute of Neuroscience, UCLouvain, Belgium. All participants were screened for eligibility of TMS experiments following standard guidelines (Lefaucheur et al., 2011; Rossi et al., 2009), which included having metallic implants in the brain or a diagnosis of epilepsy as the main exclusion criteria. The study was approved by the Saint-Luc Hospital Ethical committee (reference number: 30LXX111372). All individuals gave written informed consent prior to their participation and were financially compensated (€15).

### Materials

#### Transcranial Magnetic Stimulation

TMS was delivered using a BiStim device (Magstim Co., UK) capable of producing single-pulse monophasic stimulation. TMS was delivered in ‘BiStim mode’ (Do et al., 2020) via a single 40mm figure-of-eight alpha coil (Magstim Co., UK). Timing and delivery of TMS was controlled using a power/micro 1401 data acquisition device via Signal software version 6.04 (both by Cambridge Electronic Design, UK). A custom-made serial port interface cable (Cambridge Electronic Design, 2012) enabled direct control of the stimulator using the data acquisition software/hardware. Neuronavigation (Visor2, ANT Neuro, The Netherlands) was used to identify and maintain correct coil position.

#### Electromyography (EMG)

Skin surface EMG was recorded from the First Dorsal Interosseus (FDI) muscle using a D360 amplifier (Digitimer Ltd., Hertfordshire, UK). The skin was cleaned with alcohol before electrode placement. EMG was acquired with two circular Ag/AgCl self-adhesive surface electrodes (9mm diameter) using a belly-tendon montage, and a ground/reference electrode over the ulnar styloid process. EMG signals were amplified with a gain of 1000, online bandpass (10Hz-500Hz) and Notch-filtered (50Hz), and digitized at 4KHz using a 1401 unit.

### General Procedures

Participants sat comfortably with their dominant hand resting on a table. They were asked to stay relaxed throughout the procedure and to keep their eyes open looking at a blank black screen.

#### Motor Hotspot

The motor hotspot was defined as the optimal position of the coil to produce reliable MEPs in the FDI at the lowest intensity of stimulation (Rossini et al., 2015). TMS was delivered with the coil handle pointing backwards (i.e., to generate posterior-anterior currents) (Siebner et al., 2022), initially oriented 45° from the midline in the horizontal axis and placed tangentially over the scalp, with the final position adjusted individually. Starting at a low intensity (30% MSO), stimulation was increased in steps of 5% MSO while keeping the coil position approximately constant until an MEP exceeding 50 µV was observed. At that intensity, the coil was systematically moved to adjacent scalp positions to identify the site yielding the largest and most consistent responses, which was designated as the motor hotspot.

Given that the determination of the RMT depends on the exact coil placement during the measurement, a neuronavigation system was used to register the position and orientation in 3D space. This system used a standard brain template, as it was only used to record and reproduce the neurophysiologically defined hotspot (i.e., the motor hotspot based on MEP responses) throughout the session for each participant. This procedure allowed all subsequent measurements to use this position consistently. We also recorded the % MSO at which the hotspot was identified, the ‘hotspot intensity’ for later use (see below).

### Defining the Resting Motor Threshold (RMT)

Based on the description by the International Federation of Clinical Neurophysiology (IFCN) (Rossini et al., 2015), the RMT was defined as the lowest intensity in % of MSO at which MEPs with an amplitude ≥50μV could be elicited in at least 5 out of 10 trials (Rossini et al., 2015). For the purposes of estimating the RMT, a trial was therefore considered ‘valid’ if it produced an MEP of ≥50μV and was otherwise considered ‘invalid’.

### Stopping Criteria for the RMT

Based on the definition above, when assessing whether a given value of MSO could represent the RMT, it was not always necessary to collect exactly 10 trials. Instead, we stopped testing at a given intensity as soon as:

1) 5 valid trials were collected (i.e. it was now possible that this % MSO could represent the RMT)
OR
2) 6 invalid trials were collected (i.e. it was no longer possible for this % MSO to represent the RMT).

### Manual Assessments of the RMT

Stimulation began at the previously identified ‘hotspot intensity’. Responses were visually assessed using cursors positioned at ±25μV on screen. Although the steps of MSO used to find the RMT were technically left to the discretion of the experimenter, we systematically aimed to apply the same adjustment steps across participants. After testing the hotspot intensity, we typically moved with increases/decreases of 3% MSO followed by finer 1-2% steps to reach to RMT intensity (Stokes et al., 2013). To minimize potential trial-to-trial carryover effects and reduce the likelihood of participants from directly anticipating the delivery of the next pulse (Spiro et al., 2026), TMS was delivered at a random interval of 7.5±2.5 seconds (range: 5-10 seconds) controlled by the Signal software.

Two researchers jointly performed all manual determinations of the RMT. One researcher (MMV) had extensive TMS expertise (>3 years) and formal training in non-invasive brain stimulation. The other researcher (LFB) was trained extensively before starting data collection for this study, but did not receive a formal training and had no other previous hands-on experience with TMS. While one researcher was more experienced than the other, both performed the assessments jointly and reached a consensus value for each measurement. This collaborative approach mitigates the risk of individual operator error and is consistent with best practice in clinical neurophysiology settings (Fried et al., 2021).

### Automated Assessment of the RMT using the “RMT-Finder” procedure

#### General Procedure & Binary Search Algorithm

Automated assessments of the RMT were conducted using a custom written Signal script based on a binary search algorithm. The general principle of a binary search is to rapidly converge towards a target value by reducing the search space by half at each step (Nowak, 2008).

The general procedure is illustrated in Figure 1B. Automated assessments searched for the RMT based on a ‘search space’ between two values (the minimum and maximum values of MSO that could reasonably be expected to include the RMT). At each iteration of the algorithm, the current mid-point of this search space was identified, and trials were collected at the corresponding % MSO until one of the two stopping criteria was achieved. If a tested intensity gave 5 valid MEPs, the algorithm recorded it as a possible value for the RMT, then reduced the search space by discarding all higher values (i.e. the maximum search value was set to the intensity just tested −1% MSO), and the search process continued across this reduced search space. By contrast, if a tested intensity gave 6 invalid MEPs, the algorithm reduced the search space by removing that intensity and all values below it (i.e. the minimum value was set to the last intensity examined +1% MSO). This process was repeated iteratively until the search space was exhausted; at this point the lowest intensity examined that gave at least 5 valid MEPs was determined as the RMT. A key feature of this process is that it always confirms that the % MSO for the RMT gives at least 5 valid responses, and that the value 1% MSO below it does not.

**Figure 1.**
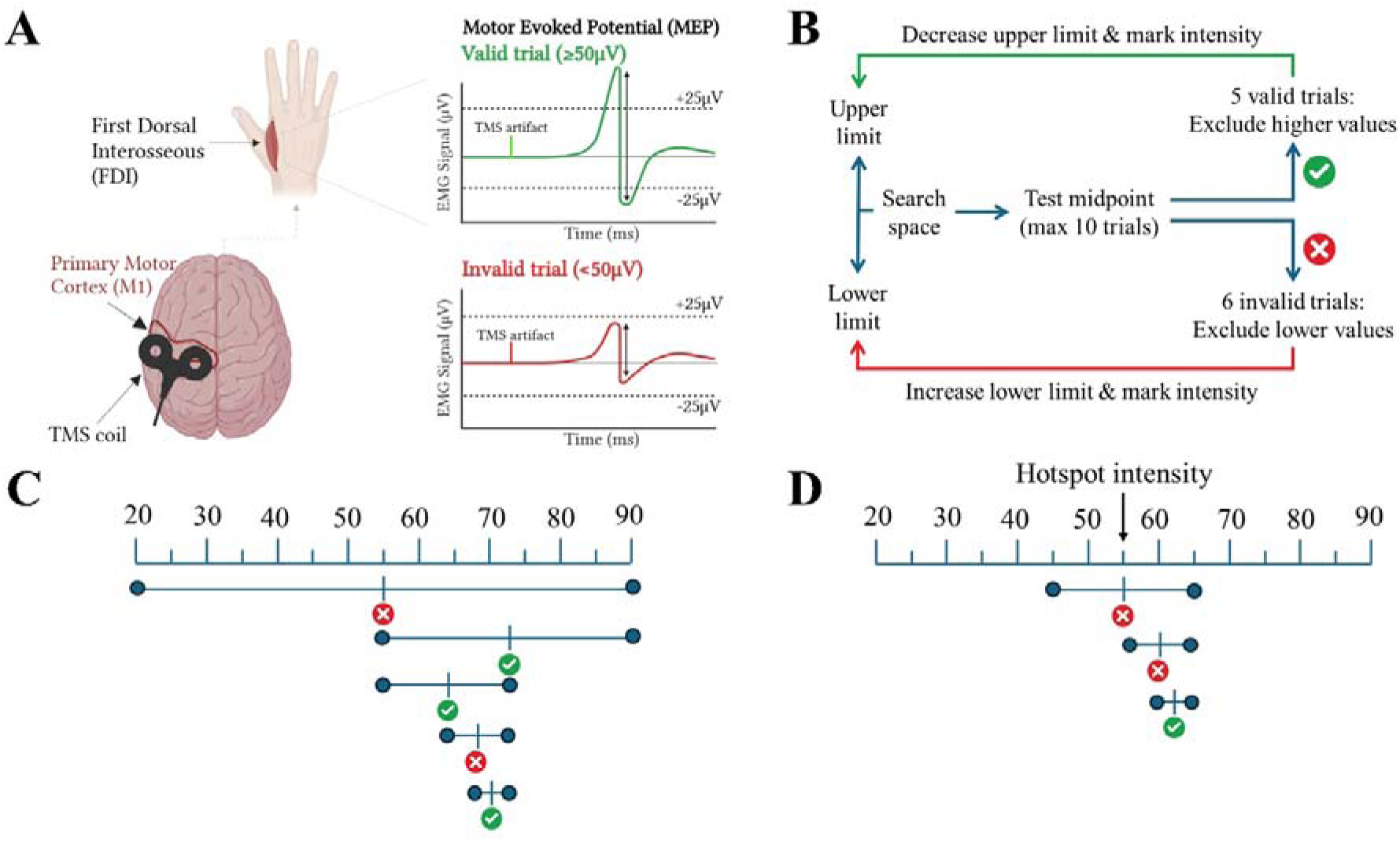
Methodological overview. **A)** Schematic representation of the participant set-up. Motor Evoked Potentials (MEPs) are measured from the First Dorsal Interosseous (FDI) of the dominant hand using Transcranial Magnetic Stimulation (TMS) applied to the Primary Motor Cortex (M1). Trials with a MEP with ≥50μV peak-to-peak amplitude are considered ‘valid’ trials, and <50 μV ‘invalid’. **B)** Visual representation of the iterative search procedure. The algorithm begins at the midpoint between predefined lower and upper intensity limits and attempts to identify at least 5 valid trials, until either 5 valid or 6 invalid trials are collected (i.e. a maximum of 10 trials). Based on the outcomes, it updates either the lower or upper boundary, progressively narrowing the search space. The process continues until the lower and upper limits converge. At this point, the algorithm selects the lowest intensity that provided 5 valid responses, which is defined as the Resting Motor Threshold (RMT), expressed in terms of % of Maximal Stimulator Output (% MSO). **C)** Illustration of the stepwise intensity adjustment performed by the original “Auto” method. Stimulation intensities tested during the procedure are marked by the symbol “⎮”. The circles indicate the (respective) current lower and upper boundaries of the search space, which is reduced by half at each iteration step as the algorithm converges towards the RMT. Red crosses mark excluded values, green check marks indicate included values. **D)** The same stepwise intensity adjustments process as illustrated in panel C is shown here, but using the Fast_Auto_ procedure. This version requires fewer iterations as the initial search space is limited to ±10% MSO of the hotspot intensity.

Background muscle activation can significantly increase the amplitude of MEPs. In the present study the RMT Finder algorithm provided measurements of the root mean square (RMS) EMG activity in the 100ms prior to the application of TMS (specifically between 102-2ms prior to the trigger signal in order to avoid interference from the TMS stimulus artefact), and the experimenters verified that this value did not exceed 10uV. The algorithm as provided (see “Data and Code Availability” statement) includes the possibility to automatically reject any trials where such a threshold is exceeded.

### Experiment 1: Comparing Manual and Auto RMT Assessments

#### Participants

A total of 24 participants completed the study (age = 27.4±8.6 years (mean±SD), 12 females, 12 males, 23 right-handed, 1 left-handed).

#### Procedure

The goal of Experiment 1 was to compare whether our automated RMT-Finder algorithm (‘Auto’) produced similar values as the Manual method, and to compare the test-retest reliability of the Automated and Manual methods. To examine their convergence, the initial search space of the Auto procedure was set between 20% and 90% of MSO, as previous data indicated that the RMT would be unlikely to be observed beyond this range when using a BiStim configuration (Do et al., 2020). Using this search space, the algorithm would always begin by first testing 55% of MSO (the mid-point of the pre-defined range; see Figure 2B) and would reach the final value for the RMT in a maximum of 7 iterations. To maintain consistency across manual and automated methods, the inter-trial interval was equivalent to the manual method (7.5 ± 2.5 seconds).

**Figure 2.**
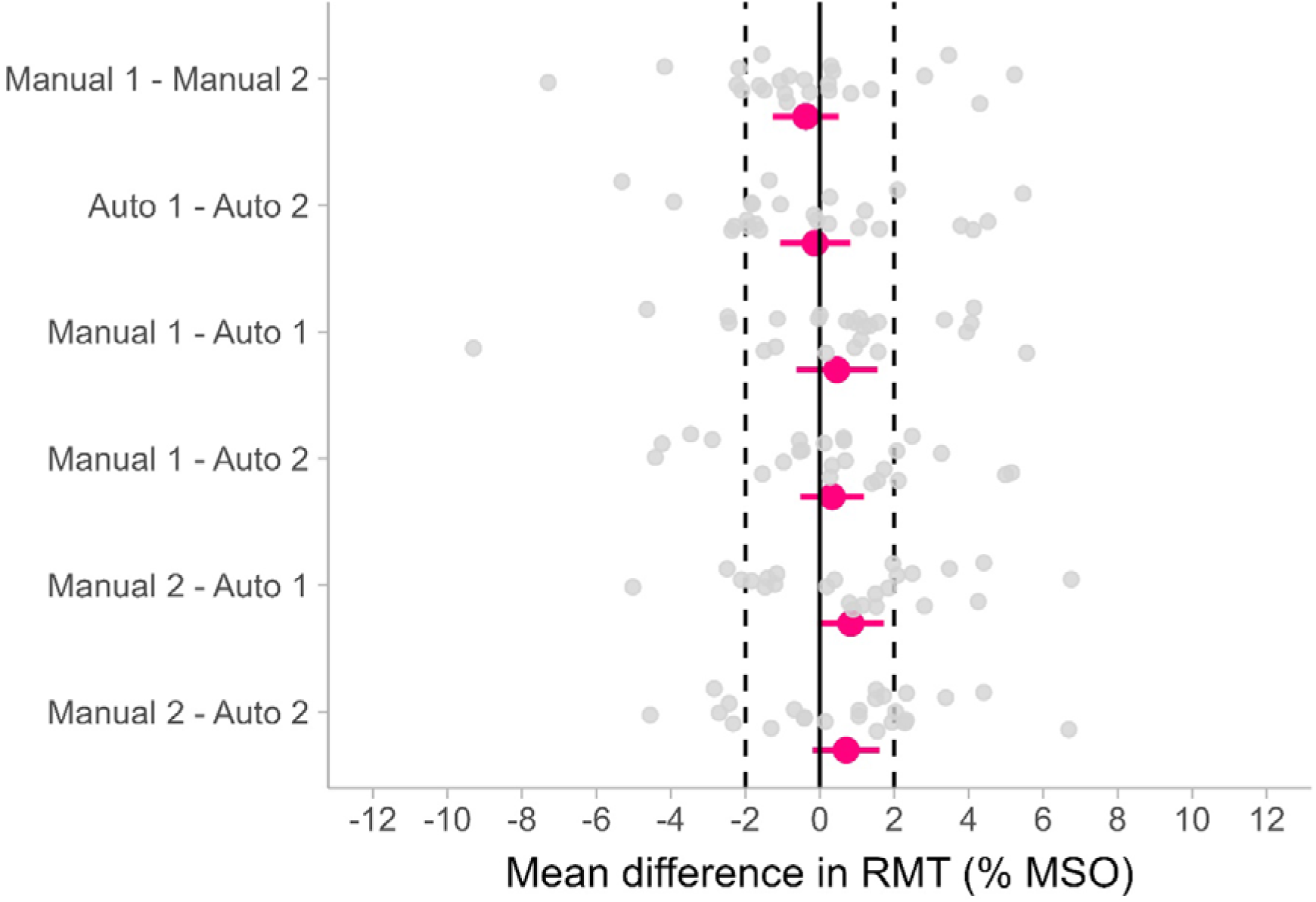
Equivalence testing for Manual and Automated methods to determine the Resting Motor Threshold (RMT) in Experiment 1 (N=24). Each colored point represents the mean difference between two measurements. Error bars indicate the 90% confidence interval. Grey dots present the data for individual participants. The dashed vertical lines mark the predefined equivalence bounds in % of Maximal Stimulator Output (% MSO).

A within-participants design was used given the substantial inter-individual variability of the RMT (Chagas et al., 2018). For each participant the RMT was measured twice using the Manual method and twice using the Auto method, with at least one minute interval between each measurement. This double assessment was designed to evaluate both the repeatability and the variability of each method independently, as well as the agreement between the two methods. The order of the four measurements was randomized (counterbalanced across participants) to minimize potential order effects.

### Experiment 2: Developing and Validating the Fast_Auto_ RMT-Finder Procedure

#### Participants

A total of 24 participants completed the study (mean age = 25.8±6.6 years (mean±SD), 16 females, 8 males, 22 right-handed, 2 left-handed).

#### Procedure

Experiment 2 aimed to optimize the practical application of the RMT-Finder procedure by making it faster while maintaining reliability. This ‘Fast_Auto_’ version of the automated procedure involved several modifications. First the search space was reduced and defined by taking a range based on the hotspot intensity ±10% MSO. This provides a range and starting point that should be closer to the RMT of the individual participant and requires only 4 to 5 iterations to determine the RMT. To accelerate the procedure even further, a fixed inter-trial interval of 4.5 seconds was used to accelerate the procedure even further; this was chosen based on the minimum time the stimulator requires to re-arm between pulses, and was longer than the minimum recommended inter-trial-interval of 4s (Osnabruegge et al., 2023).

The RMT was measured once manually, once with the previous version (‘Auto’) of the RMT-Finder procedure, and twice with the new (‘Fast_Auto_’) version, with at least one minute interval between each measurement. This was done to test the repeatability of the Fast_Auto_ procedure, as well as its equivalence with the Manual and Auto methods. The order of the different measurements was randomized (counterbalanced across participants) to reduce potential order effects.

Experiment 2 also implemented a blinding procedure to reduce the risk of experimenter bias in manual measurements. Specific roles were assigned to two experimenters at the beginning of the experiment. Experimenter 1 was responsible for applying TMS; they identified the motor hotspot and held the coil over this position throughout the rest of the testing session. Experimenter 2 was responsible for managing stimulation intensity (including during hotspot identification) and for loading the files and configurations into the Signal software. This procedure effectively allowed Experimenter 1 to be blind to the exact intensity (% MSO) of TMS being delivered during the experiment. During hotspotting and manual RMT assessment, Experimenter 1 was responsible for reviewing MEPs and determining if they were ‘valid’ or ‘invalid’; they then asked Experimenter 2 to increase or decrease the intensity of TMS according to the responses they observed, without knowing the specific values of % MSO being applied.

We also measured the time required to identify the RMT for each technique, taken from the first to the last pulse of TMS required. The duration of manual measurements was taken with a stopwatch by Experimenter 2, while the duration of automated measurements was computed by the automated script.

### Data analysis

All statistical analyses were conducted using R version 4.5.1 (R Core Team 2025). In each experiment we conducted planned pairwise comparisons for each measurement taken with each approach to finding the RMT.

#### Reliability & Agreement

The Intraclass Correlation Coefficient (ICC) with a two-way mixed-effects model for single measures and absolute agreement (2,1) was used (Koo and Li, 2016). This coefficient ranges between 0-1, higher values indicating greater reliability. We also analysed measurement error via the Standard Error of Measurement (SEM), calculated as SEM = SD * √(1 – ICC), where SD is the pooled standard deviation for the pair of measurements and the ICC is obtained from the previous step (Denegar and Ball, 1993). Limits of agreement were visualized by Bland-Altman plots (see Supplementary Materials) (Bland and Altman, n.d.).

#### Equivalence

The Two One-Sided Test (TOST) procedure was used. This approach allows testing whether the difference between two conditions falls within a predefined equivalence margin. Any differences between conditions within that region can be considered practically negligible (i.e. similar to 0) (Lakens, 2017). We a priori defined the desired margin as ranging ±2% MSO (Badran et al., 2019) (we note that while previous studies have not performed direct equivalence tests, they appear to have generally accepted a range of ±5% MSO (Julkunen, 2019)). As each participant provided data from all measurements, we used paired TOSTs.

#### Efficiency

Previous studies have measured the efficiency of RMT assessment approaches based on the number of required pulses. We anticipated that an optimized automated approach could also significantly reduce the overall time required for RMT procedures (including minimizing the time required to measure each MEP, determine whether the intensity of stimulation should be modified, and physically implement changes), allowing data collection with minimal inter-trial intervals. We also note that knowing the overall time required to perform the procedure is most informative when planning experiments. In Experiment 2 we therefore measured the efficiency of each technique in terms of the number of intensities tested, the overall number of pulses and time required to determine the RMT for each approach.

#### Sample Size Estimation

Power analysis was conducted based on a separate sample of 4 participants (see Supplementary Materials). Calculations were based on paired TOST tests, with equivalence bounds set at ±2% MSO. To achieve a statistical power of 95% with alpha = 0.05, considering the standard deviation of the difference between two consecutive automated measurements (SD = 2% MSO after accounting for variance underestimation) the sample size required was 13 pairs of data. To account for possible dropouts, further underestimations of the variance, and to allow a fully counterbalanced design, we collected a final sample of 24 participants in each experiment.

## Results

### Experiment 1

#### Descriptive Statistics

Both the manual and automated methods yielded qualitatively similar RMTs (Mean±SD, Manual 1: 58.8±10.9% MSO, Manual 2: 58.9±11.2% MSO, Auto 1: 58.4±10.3% MSO, Auto 2: 58.6±10.6% MSO; Supplementary Figure 1).

#### Reliability

All ICCs were ≥0.96 (Table 1), reflecting excellent reliability both within and between methods. Bland-Altman plots (see Supplementary Figure 2), indicated minimal bias and narrow limits of agreement, illustrating high consistency within and across methods.

**Table 1.** Reliability of Resting Motor Threshold (RMT) measurements across methods in Experiment 1 (N=24).

| Comparison | ICC | 95% CI | SEM |
| --- | --- | --- | --- |
| Manual 1 vs Manual 2 | 0.98 | [0.94, 0.99] | 1.77 |
| Auto 1 vs Auto 2 | 0.97 | [0.93, 0.99] | 1.87 |
| Manual 1 vs Auto 1 | 0.96 | [0.91, 0.98] | 2.17 |
| Manual 1 vs Auto 2 | 0.98 | [0.95, 0.99] | 1.71 |
| Manual 2 vs Auto 1 | 0.97 | [0.94, 0.99] | 1.85 |
| Manual 2 vs Auto 2 | 0.97 | [0.94, 0.99] | 1.85 |

Abbreviations: CI = Confidence Interval; ICC = Intraclass Correlation Coefficient; SEM = Standard Error of Measurement.

#### Equivalence

Pairwise comparisons conducted using TOST analysis indicated that all measurements gave equivalent results within ±2% MSO (see Figure 2, Table 2).

**Table 2.** Results of equivalence tests via paired Two One-Sided Test (TOST) procedure for all comparisons between the different Resting Motor Threshold (RMT) assessment methods in Experiment 1 (N=24). The table shows the results of the TOST used to assess equivalence. The lower bound test evaluates the null hypothesis that the true difference is ≤ −2 %MSO, and the upper bound test evaluates the null hypothesis that the true difference is ≥ +2 %MSO. When both one-sided tests yield p < .05, the null hypotheses of non-equivalence are rejected, and the TOST procedure provides evidence that the true difference between methods falls within the equivalence region of [−2, +2] %MSO.

| Comparison | Lower bound | Upper bound |
| --- | --- | --- |
| Manual 1 vs Manual 2 | $t_{23}=3.14, p=0.002$ | $t_{23}=-4.59, p<0.0001$ |
| Auto 1 vs Auto 2 | $t_{23}=3.41, p=0.001$ | $t_{23}=-3.87, p=0.0003$ |
| Manual 1 vs Auto 1 | $t_{23}=3.89, p=0.0004$ | $t_{23}=-2.44, p=0.011$ |
| Manual 1 vs Auto 2 | $t_{23}=4.68, p<0.0001$ | $t_{23}=-3.34, p=0.001$ |
| Manual 2 vs Auto 1 | $t_{23}=5.48, p<0.0001$ | $t_{23}=-2.26, p=0.017$ |
| Manual 2 vs Auto 2 | $t_{23}=5.14, p<0.0001$ | $t_{23}=-2.45, p=0.011$ |

### Experiment 2

#### Descriptive Statistics

As in Experiment 1, all methods provided qualitatively similar estimates of the RMT with comparable ranges (Mean±SD, Manual: 59.1±11.2% MSO, Auto: 59.7±12.0% MSO, Fast_Auto_ 1: 59.6±10.8% MSO, Fast_Auto_ 2: 60.5±11.4% MSO; Supplementary Figure 3).

#### Reliability

All ICCs were ≥0.95 (Table 3), reflecting excellent reliability both within and between methods. Bland-Altman plots (Supplementary Figure 3), indicated minimal bias and narrow limits of agreement, illustrating high consistency within and across methods.

**Table 3.**
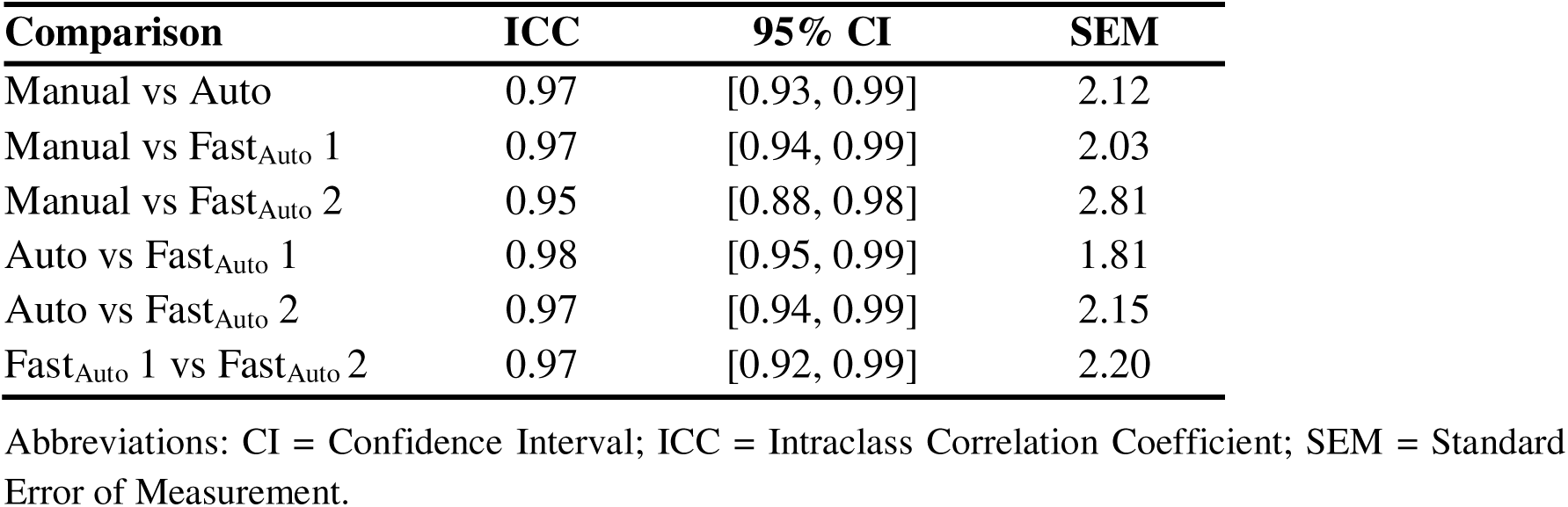
Reliability of Resting Motor Threshold (RMT) measurements across methods in Experiment 2 (N=24).

#### Equivalence

Pairwise comparisons conducted using TOST analysis indicated that the majority of comparisons gave equivalent results within ±2% MSO (Figure 3, Table 4). A single comparison (Manual - Fast_Auto_ 2) slightly exceeded these bounds but remained equivalent within ±3% MSO.

**Figure 3.**
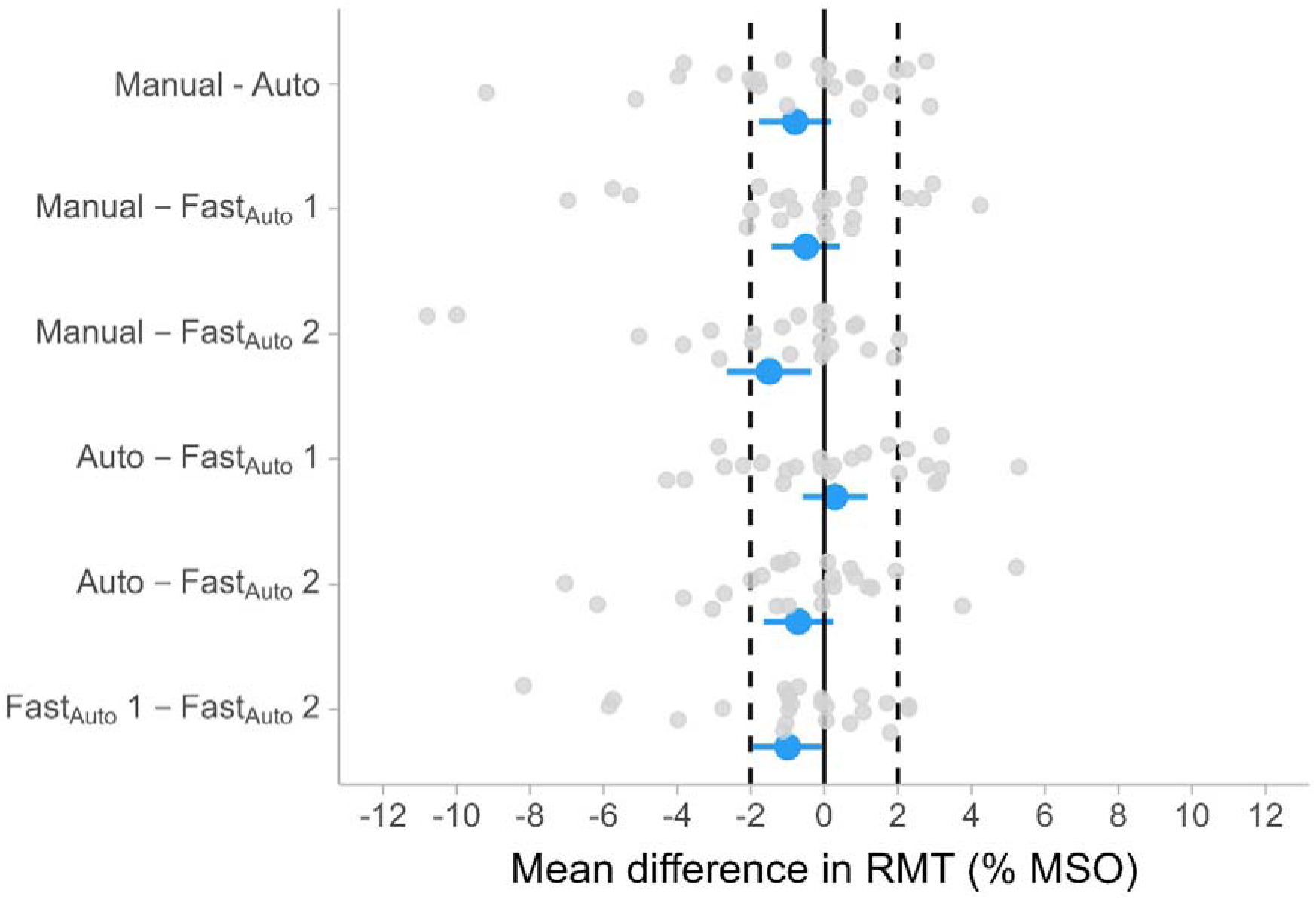
Equivalence testing for Manual and Automated methods to determine the Resting Motor Threshold (RMT) in Experiment 2 (N=24). Each colored point represents the mean difference between two measurements. Error bars indicate the 90% confidence interval. Grey dots present the data for individual participants. The dashed vertical lines mark the predefined equivalence bounds in % of Maximal Stimulator Output (% MSO).

**Table 4.** Results of equivalence tests via paired Two One-Sided Test (TOST) procedure for all comparisons between manual and automated Resting Motor Threshold (RMT) determination methods in Experiment 2 (N=24). The table shows the results of the TOST used to assess equivalence. The lower bound test evaluates the null hypothesis that the true difference is ≤ −2 %MSO, and the upper bound test evaluates the null hypothesis that the true difference is ≥ +2 %MSO. When both one-sided tests yield p < .05, the null hypotheses of non-equivalence are rejected, and the TOST procedure provides evidence that the true difference between methods falls within the equivalence region of [−2, +2] %MSO

| Comparison | Lower bound | Upper bound |
| --- | --- | --- |
| Manual vs Auto | $t_{23}=2.19, p=0.0019$ | $t_{23}=-5.03, p<0.0001$ |
| Manual vs Fast<sub>Auto</sub> 1 | $t_{23}=2.89, p=0.004$ | $t_{23}=-4.81, p<0.0001$ |
| Manual vs Fast<sub>Auto</sub> 2 | $t_{23}=0.79, p=0.22$ | $t_{23}=-5.44, p<0.0001$ |
| Auto vs Fast<sub>Auto</sub> 1 | $t_{23}=4.68, p<0.0001$ | $t_{23}=-3.51, p=0.001$ |
| Auto vs Fast<sub>Auto</sub> 2 | $t_{23}=2.43, p=0.011$ | $t_{23}=-5.11, p<0.0001$ |
| Fast<sub>Auto</sub> 1 vs Fast<sub>Auto</sub> 2 | $t_{23}=1.92, p=0.033$ | $t_{23}=-5.73, p<0.0001$ |

#### Efficiency

Efficiency data is summarized in Table 5. Both the Manual and Fast_Auto_ procedures required an average of 33-34 pulses to identify the RMT. The original Auto procedure required more pulses than the other procedures, presumably due to its wide initial search range. By contrast, while both the Manual and original Auto procedure required an average of around 5 minutes to identify the RMT, the Fast_Auto_ procedure required approximately half this time, typically identifying the RMT in <3 minutes.

**Table 5.**
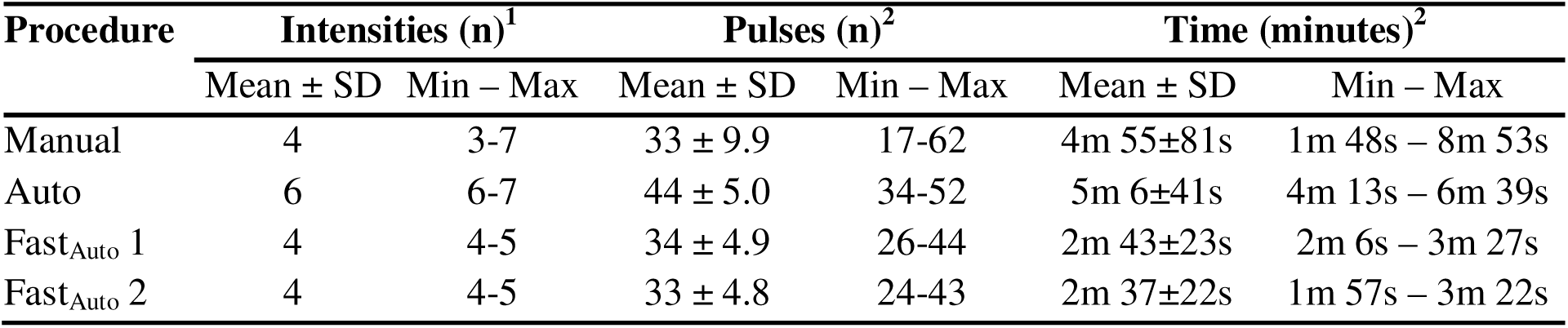
Efficiency measures in Experiment 2 (N=24), illustrating the number of intensities tested, the overall number pulses, and the time required to find the Resting Motor Threshold (RMT) for each procedure.

| Procedure | Intensities (n)<sup>1</sup> |  | Pulses (n)<sup>2</sup> |  | Time (minutes)<sup>2</sup> |  |
| --- | --- | --- | --- | --- | --- | --- |
|  | Mean ± SD | Min – Max | Mean ± SD | Min – Max | Mean ± SD | Min – Max |
| Manual | 4 | 3-7 | 33 ± 9.9 | 17-62 | 4m 55±81s | 1m 48s – 8m 53s |
| Auto | 6 | 6-7 | 44 ± 5.0 | 34-52 | 5m 6±41s | 4m 13s – 6m 39s |
| Fast<sub>Auto</sub> 1 | 4 | 4-5 | 34 ± 4.9 | 26-44 | 2m 43±23s | 2m 6s – 3m 27s |
| Fast<sub>Auto</sub> 2 | 4 | 4-5 | 33 ± 4.8 | 24-43 | 2m 37±22s | 1m 57s – 3m 22s |

## Discussion

The present manuscript introduces an automated relative-frequency assessment approach that can accurately identify the RMT in, on average, <3 minutes. Across two experiments the RMT-Finder method demonstrated strong reliability, agreement, and equivalence with traditional manual procedures. Conducting multiple measurements with each version of the algorithm confirmed that the RMT-Finder procedure gave consistent results.

Across both experiments, each version of the RMT-Finder procedure demonstrated excellent agreement with manual measurements (all ICCs ≥0.95). Moreover, repeated assessments of the RMT using the Fast_Auto_ procedure also yielded excellent test-retest reliability (ICC=0.97). These results are similar to previous studies that have compared within- and between-technique approaches to identifying the RMT (Ah Sen et al., 2017; Silbert et al., 2013). While we were unable to find previous studies that performed direct equivalence tests between their procedures and manual methods, previous studies suggest their RMT determination procedures agree with manual methods within ±5% MSO (Julkunen, 2019). To the best of our knowledge the present study is therefore the first to demonstrate statistically equivalent procedures; all comparisons between our automated and manual measurements were statistically equivalent to within ±3% MSO, and all but one comparison were statistically equivalent within ±2% MSO. Given that ICC measurements do not always consider absolute agreement between measurements, we propose equivalence testing procedures may provide a clearer approach to demonstrate two procedures provide similar measurements. Overall, these results demonstrate the success of our procedure, given that the RMT-Finder approach aimed to replicate and automate several of the steps involved in taking traditional manual relative-frequency-based measurements of the RMT.

Results of Experiment 2 demonstrate that the revised Fast_Auto_ version of the RMT-Finder procedure was rapidly able to identify the RMT while maintaining test-retest reliability. Previous attempts to optimize the efficiency of identifying the RMT have generally focused on the number of pulses required, with 30 pulses being recommended for curve-fitting approaches (Koponen and Peterchev, 2022). We note this is comparable to the average of 33-34 pulses taken by the present Fast_Auto_ procedure, which requires fewer pulses than approaches based on current IFCN guidelines. This is mainly because it reduces the search space based on the intensity used during hotspotting and therefore takes individuals’ responses into account instead of using fixed rules (e.g. starting at 50% MSO) (Mishory et al., 2004; Rossini et al., 2015). However, given the inherent trial-to-trial variability of MEP amplitudes, recent literature has argued that reliable measurements depend on collecting several trials at a given intensity (Ammann et al., 2020; Biabani et al., 2018; Chang et al., 2016). Thus, rather than focusing on reducing the overall number of pulses, it may be more appropriate to collect multiple trials per intensity and simultaneously minimize the time this takes. The present Fast_Auto_ procedure was able to accurately determine the RMT in <3 minutes on average, making it not only a useful tool to automate the process, but also a procedure directly relevant for experiments or clinical assessments where faster measurements are desirable.

Beyond its relative speed and efficiency, the automated nature of the RMT-Finder procedure means that it also places minimal burden on the experimenter(s) using it. Previous approaches to identifying the RMT either effectively require a single experimenter to multi-task (i.e. switch attention between maintaining optimal coil position and assessing the amplitude of MEP responses, remembering previous results of trials at a given intensity, and making/executing decisions about modifying the stimulator output) or split these tasks between different experimenters (Groppa et al., 2012). Both these options introduce the possibility for errors. By contrast, it is completely feasible for a single experimenter to conduct the full RMT-Finder procedure alone, allowing them to devote their full attention to maintaining the position of the coil. Moreover, the search-algorithm (i.e. search space, step size, stopping criteria, etc) can be fully defined, improving the direct replicability of this procedure between laboratories.

We note the current implementation was validated only in healthy participants targeting a single muscle representation, which may limit generalization. The present version of the algorithm is also limited to use with CED’s Signal software to measure MEP amplitudes and communicate with the TMS device; however, it is feasible to extend the procedures outlined in this study to other applications such as open-source software (e.g. MagPy (McNair, 2017)). These limitations can be readily addressed in future work.

While the present findings support the use of the automated RMT finder across a broad range of applications, we wish to emphasise that automated tools of this kind are best regarded as an aid to, rather than a replacement for, operator expertise. RMT determination is identified by the IFCN training guidelines (Fried et al., 2021) as a core foundational skill for all TMS practitioners, and the process of finding RMT manually cultivates an understanding of coil handling, MEP variability, and motor cortex excitability that is difficult to acquire through automated methods alone.

## Conclusion

The present manuscript developed an automated procedure to determine the RMT. Results were in strong agreement with the RMT as identified using manual approaches (ICCs≥0.95), and all results were equivalent to within ±3% MSO, with the vast majority within ±2% MSO. It also showed similar test-retest reliability, while substantially reducing the amount of time required compared to manual measurements. This automated procedure is also more convenient for the experimenter, allowing them to focus fully on maintaining optimal coil positioning. These advantages represent a meaningful step forward in optimizing TMS protocols, particularly in settings where time efficiency and procedural standardization are crucial, such as in clinical applications.

## Acknowledgements

We would like to thank Prof. Philippe Lefèvre, Prof. André Moreaux and Prof. Julie Duqué for providing equipment necessary to conduct this study.

## Data & Code availability

The data and code used for analysis in this study can be found in the corresponding OSF repository (https://osf.io/4cq92/overview). The latest version of the materials to use the ‘RMT-Finder’ can be found in its dedicated Github repository (https://github.com/mmorenoverdu/RMT-Finder).

## Author Roles (CRediT)

| Author | Conceptualization | Methodology | Software | Validation | Formal Analysis | Investigation | Resources | Data Curation | Writing – original draft | Writing – reviewing & editing | Visualization | Supervision | Project Administration | Funding Acquisition |
| --- | --- | --- | --- | --- | --- | --- | --- | --- | --- | --- | --- | --- | --- | --- |
| LFB | X | X |  | X | X | X |  | X | X | X | X |  |  |  |
| MMV | X | X |  | X | X | X |  | X | X | X | X | X |  | X |
| BMW |  |  |  |  |  | X |  |  |  | X |  |  |  |  |
| JJL |  |  |  |  |  |  | X |  |  | X |  |  |  |  |
| EVC |  |  |  |  |  |  |  |  |  | X |  |  |  |  |
| CT |  |  |  |  |  |  |  |  |  | X |  |  |  |  |
| RMH | X | X | X |  |  |  | X |  | X | X |  | X | X | X |

## References

Ah Sen CB, Fassett HJ, El-Sayes J, Turco CV, Hameer MM, Nelson AJ. Active and resting motor threshold are efficiently obtained with adaptive threshold hunting. PLoS ONE 2017;12:e0186007. 10.1371/journal.pone.0186007.

Ammann C, Guida P, Caballero-Insaurriaga J, Pineda-Pardo JA, Oliviero A, Foffani G. A framework to assess the impact of number of trials on the amplitude of motor evoked potentials. Sci Rep 2020;10:21422. 10.1038/s41598-020-77383-6.

Awiszus F. Chapter 2 TMS and threshold hunting. Supplements to Clinical Neurophysiology, vol. 56, Elsevier; 2003, p. 13–23. 10.1016/S1567-424X(09)70205-3.

Awiszus F, Borckardt, Jeffrey J. TMS motor threshold assessment tool (MTAT 2.0) 2011.

Badran BW, Ly M, DeVries WH, Glusman CE, Willis A, Pridmore S, et al. Are EMG and visual observation comparable in determining resting motor threshold? A reexamination after twenty years. Brain Stimulation 2019;12:364–6. 10.1016/j.brs.2018.11.003.

Biabani M, Farrell M, Zoghi M, Egan G, Jaberzadeh S. The minimal number of TMS trials required for the reliable assessment of corticospinal excitability, short interval intracortical inhibition, and intracortical facilitation. Neuroscience Letters 2018;674:94–100. 10.1016/j.neulet.2018.03.026.

Bland JM, Altman DG. Measuring agreement in method comparison studies n.d.

Cambridge Electronic Design L. TMS and Magstim control using Signal: A guide to setting CED acquisition and analysis systems for TMS Studies 2012.

Chagas AP, Monteiro M, Mazer V, Baltar A, Marques D, Carneiro M, et al. Cortical excitability variability: Insights into biological and behavioral characteristics of healthy individuals. Journal of the Neurological Sciences 2018;390:172–7. 10.1016/j.jns.2018.04.036.

Chang WH, Fried PJ, Saxena S, Jannati A, Gomes-Osman J, Kim Y-H, et al. Optimal number of pulses as outcome measures of neuronavigated transcranial magnetic stimulation. Clinical Neurophysiology 2016;127:2892–7. 10.1016/j.clinph.2016.04.001.

Denegar CR, Ball DW. Assessing Reliability and Precision of Measurement: An Introduction to Intraclass Correlation and Standard Error of Measurement. Journal of Sport Rehabilitation 1993;2:35–42. 10.1123/jsr.2.1.35.

Do M, Clark GM, Fuelscher I, Kirkovski M, Cerins A, Corp DT, et al. Magstim 2002 and Bistim Mode maximum stimulus output values are not equivalent: Configuration selection is critical. Brain Stimulation 2020;13:444–6. 10.1016/j.brs.2019.12.009.

Fried PJ, Santarnecchi E, Antal A, Bartres-Faz D, Bestmann S, Carpenter LL, et al. Training in the practice of noninvasive brain stimulation: Recommendations from an IFCN committee. Clinical Neurophysiology 2021;132:819–37. 10.1016/j.clinph.2020.11.018.

Groppa S, Oliviero A, Eisen A, Quartarone A, Cohen LG, Mall V, et al. A practical guide to diagnostic transcranial magnetic stimulation: Report of an IFCN committee. Clinical Neurophysiology 2012;123:858–82. 10.1016/j.clinph.2012.01.010.

Hallett M. Transcranial Magnetic Stimulation: A Primer. Neuron 2007;55:187–99. 10.1016/j.neuron.2007.06.026.

Hamoline G, Van Caenegem EE, Waltzing BM, Vassiliadis P, Derosiere G, Duque J, et al. Accelerometry as a tool for measuring the effects of transcranial magnetic stimulation. Journal of Neuroscience Methods 2024;405:110107. 10.1016/j.jneumeth.2024.110107.

Julkunen P. Mobile Application for Adaptive Threshold Hunting in Transcranial Magnetic Stimulation. IEEE Trans Neural Syst Rehabil Eng 2019;27:1504–10. 10.1109/TNSRE.2019.2925904.

Koo TK, Li MY. A Guideline of Selecting and Reporting Intraclass Correlation Coefficients for Reliability Research. Journal of Chiropractic Medicine 2016;15:155–63. 10.1016/j.jcm.2016.02.012.

Koponen LM, Peterchev AV. Preventing misestimation of transcranial magnetic stimulation motor threshold with MTAT 2.0. Brain Stimulation 2022;15:1073–6. 10.1016/j.brs.2022.07.057.

Lakens D. Equivalence Tests: A Practical Primer for *t* Tests, Correlations, and Meta-Analyses. Social Psychological and Personality Science 2017;8:355–62. 10.1177/1948550617697177.

Lefaucheur J-P, André-Obadia N, Poulet E, Devanne H, Haffen E, Londero A, et al. Recommandations françaises sur l’utilisation de la stimulation magnétique transcrânienne répétitive (rTMS)L: règles de sécurité et indications thérapeutiques. Neurophysiologie Clinique/Clinical Neurophysiology 2011;41:221–95. 10.1016/j.neucli.2011.10.062.

McNair NA. MagPy: A Python toolbox for controlling Magstim transcranial magnetic stimulators. Journal of Neuroscience Methods 2017;276:33–7. 10.1016/j.jneumeth.2016.11.006.

Mishory A, Molnar C, Koola J, Li X, Kozel FA, Myrick H, et al. The Maximum-likelihood Strategy for Determining Transcranial Magnetic Stimulation Motor Threshold, Using Parameter Estimation by Sequential Testing Is Faster Than Conventional Methods With Similar Precision: The Journal of ECT 2004;20:160–5. 10.1097/00124509-200409000-00007.

Nowak R. Generalized binary search. 2008 46th Annual Allerton Conference on Communication, Control, and Computing, Monticello, IL, USA: IEEE; 2008, p. 568–74. 10.1109/ALLERTON.2008.4797609.

Osnabruegge M, Kanig C, Schwitzgebel F, Litschel K, Seiberl W, Mack W, et al. On the reliability of motor evoked potentials in hand muscles of healthy adults: a systematic review. Front Hum Neurosci 2023;17:1237712. 10.3389/fnhum.2023.1237712.

Rossi S, Hallett M, Rossini PM, Pascual-Leone A. Safety, ethical considerations, and application guidelines for the use of transcranial magnetic stimulation in clinical practice and research. Clinical Neurophysiology 2009;120:2008–39. 10.1016/j.clinph.2009.08.016.

Rossini PM, Barker AT, Berardelli A, Caramia MD, Caruso G, Cracco RQ, et al. Non-invasive electrical and magnetic stimulation of the brain, spinal cord and roots: basic principles and procedures for routine clinical application. Report of an IFCN committee. Electroencephalography and Clinical Neurophysiology 1994;91:79–92. 10.1016/0013-4694(94)90029-9.

Rossini PM, Burke D, Chen R, Cohen LG, Daskalakis Z, Di Iorio R, et al. Non-invasive electrical and magnetic stimulation of the brain, spinal cord, roots and peripheral nerves: Basic principles and procedures for routine clinical and research application. An updated report from an I.F.C.N. Committee. Clinical Neurophysiology 2015;126:1071–107. 10.1016/j.clinph.2015.02.001.

Siebner HR, Funke K, Aberra AS, Antal A, Bestmann S, Chen R, et al. Transcranial magnetic stimulation of the brain: What is stimulated? – A consensus and critical position paper. Clinical Neurophysiology 2022;140:59–97. 10.1016/j.clinph.2022.04.022.

Silbert BI, Patterson HI, Pevcic DD, Windnagel KA, Thickbroom GW. A comparison of relative-frequency and threshold-hunting methods to determine stimulus intensity in transcranial magnetic stimulation. Clinical Neurophysiology 2013;124:708–12. 10.1016/j.clinph.2012.09.018.

Spiro E, Zhu L, Huang Z, Ikramuddin S, Peterchev AV, Charalambous CC, et al. Inter-pulse interval and motor evoked potential variability: Bridging insights from healthy adults to post-stroke TMS protocols. Brain Stimulation 2026;19:102987. 10.1016/j.brs.2025.102987.

Stokes MG, Barker AT, Dervinis M, Verbruggen F, Maizey L, Adams RC, et al. Biophysical determinants of transcranial magnetic stimulation: effects of excitability and depth of targeted area. Journal of Neurophysiology 2013;109:437–44. 10.1152/jn.00510.2012.

Vassiliadis P, Derosiere G, Grandjean J, Duque J. Motor training strengthens corticospinal suppression during movement preparation. Journal of Neurophysiology 2020;124:1656–66. 10.1152/jn.00378.2020.

