## Supplementary Materials for "‘RMT-Finder’: an automated procedure to determine the Resting Motor Threshold for Transcranial Magnetic Stimulation"

**Pilot data for sample size calculation**

While developing the automated procedure to find the RMT, we tested 4 pilot participants and measured their RMT manually (once) and automatically (twice). These pilot data are shown in Supplementary Table 1. From these data, we calculated the difference between the pair of automated measurements for each participant and then calculated the mean difference of the sample and its SD in % MSO. The mean difference was 0.5 %MSO and the SD was 1 %MSO. Expectedly, at this sample size variance tends to be underestimated. Therefore, we conducted the calculations with a SD of 2 %MSO, which would account for that underestimation and would be a more realistic estimate. This SD was used to calculate sample size according to the statistical power in a paired TOST procedure to assess statistical equivalence (see main text in paper for details).

**Supplementary Table 1.** Pilot data for sample size calculation. The Resting Motor Threshold (RMT) and hotspot intensities are provided in % of Maximal Stimulator Output (MSO) units.

| Participant | Sex | Age | Hotspot intensity | RMT Manual | RMT Auto 1 | RMT Auto 2 |
| --- | --- | --- | --- | --- | --- | --- |
| 001 | M | 29 | 65 | 66 | 68 | 67 |
| 002 | M | 25 | 50 | 42 | 42 | 41 |
| 003 | F | 21 | 45 | 34 | 32 | 31 |
| 004 | M | 26 | 55 | 63 | 64 | 65 |

**Experiment 1: Descriptive statistics**

Supplementary Figure 1 shows the raw data from each RMT measurement in Experiment 1. Each dot is a single participant.

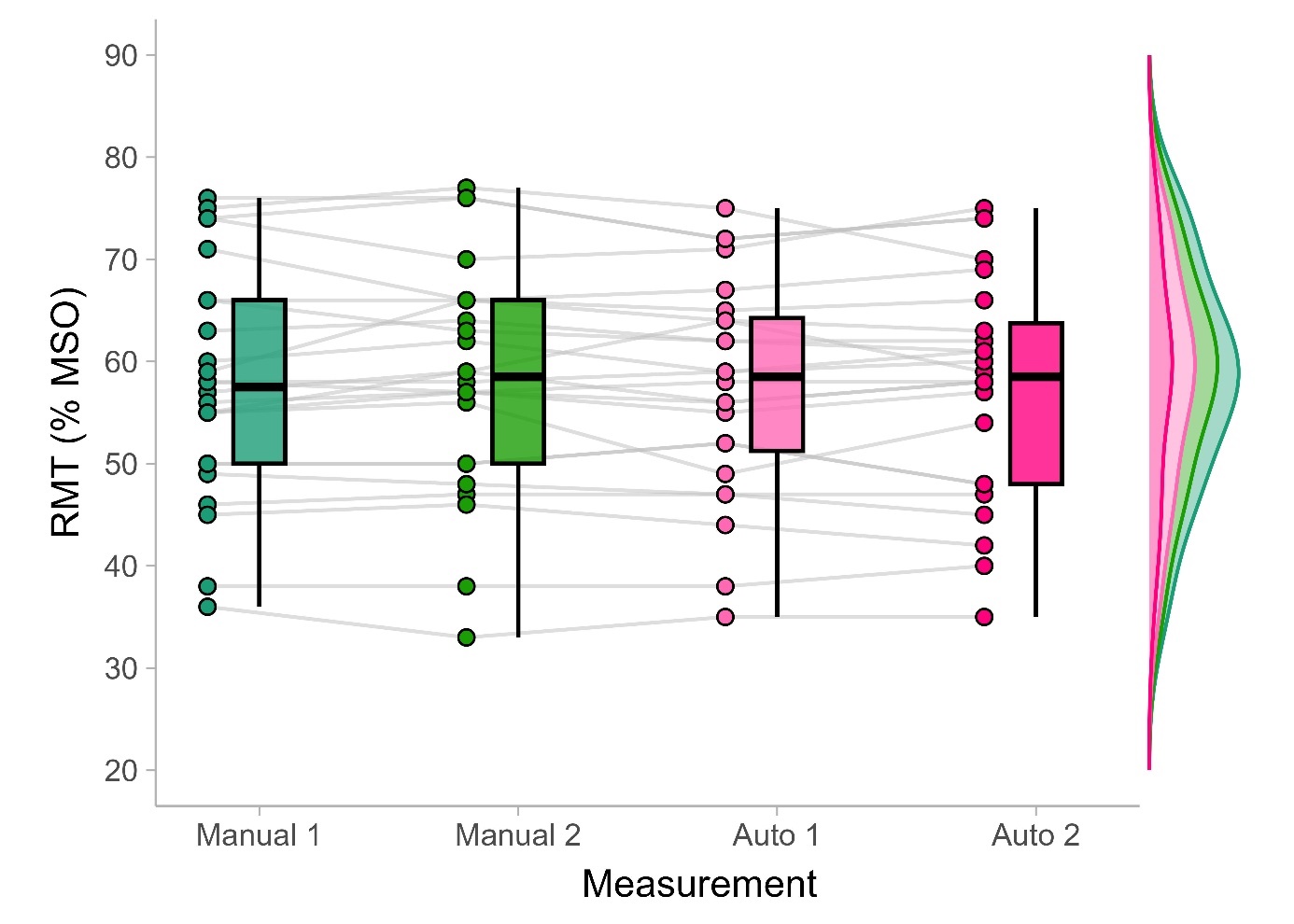

**Supplementary Figure 1.** Boxplots and distributions of the different measurements of the Resting Motor Threshold (RMT), in terms of % of Maximal Stimulator Output (% MSO), in Experiment 1 (N=24).

**Experiment 1: Limits of Agreement**

To visualize whether there are systematic biases between measurements associated with the distribution of actual values of the RMT, we plotted differences between measurements against their means for each participant through Bland-Altman plots. The plots allow the identification of the Limits of Agreement for a given pair of measurements, which reflect the level of consistency between them. The plots for Experiment 1 are shown in Supplementary Figure 2.

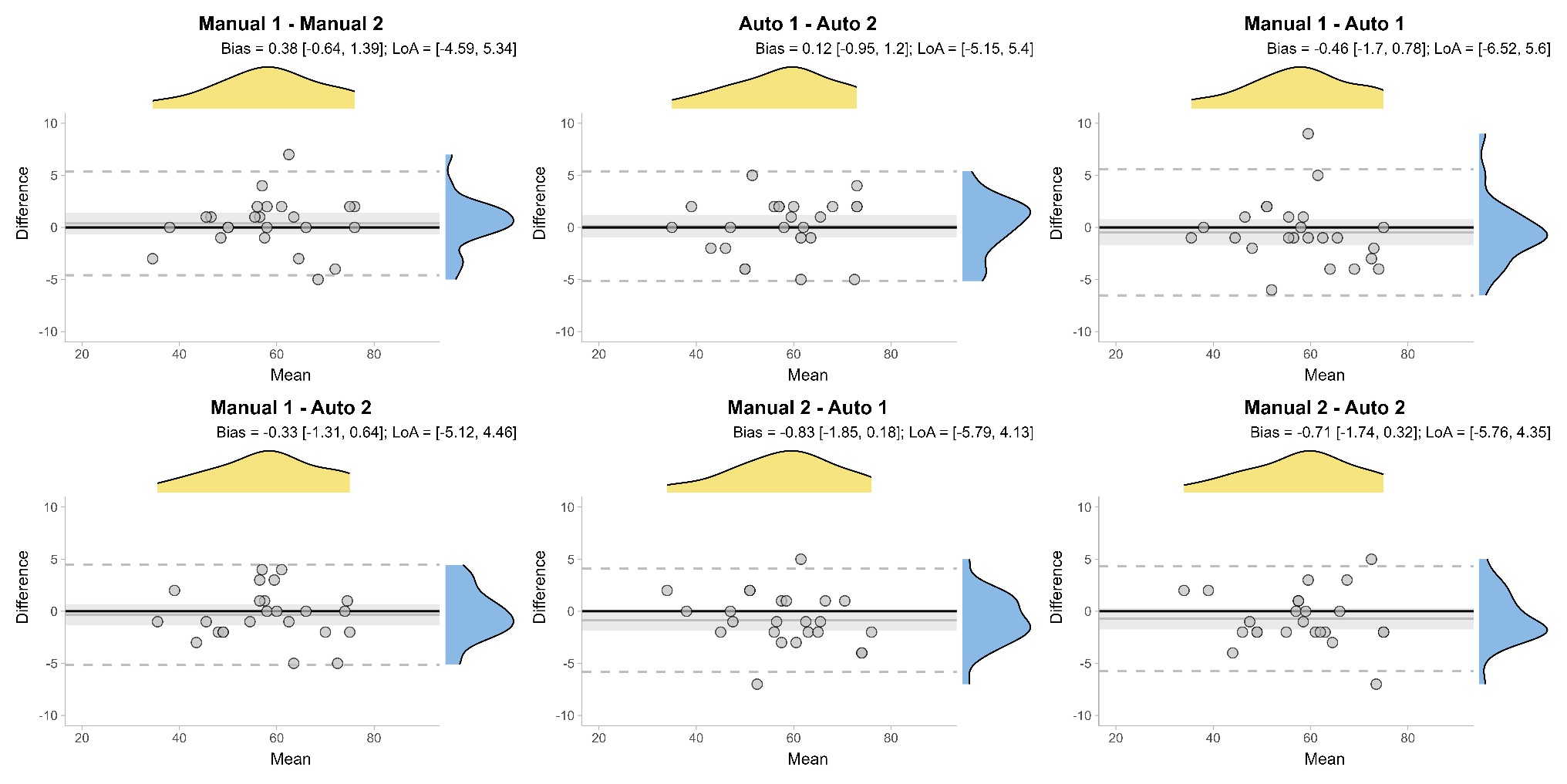

**Supplementary Figure 2.** Bland-Altman plots comparing manual and automated RMT measurements for Experiment 1. Each panel shows the mean versus the difference between two measurements. Horizontal black line shows the mean bias, and shaded areas represent the 95% confidence interval around it. Dashed horizontal lines show the Limits of Agreement (LoA). Bias and Limits of Agreement values are indicated above each panel. All comparisons show minimal bias and narrow limits of agreement.

**Experiment 2: Descriptive statistics**

Supplementary Figure 3 shows the raw data from each RMT measurement in Experiment 2.

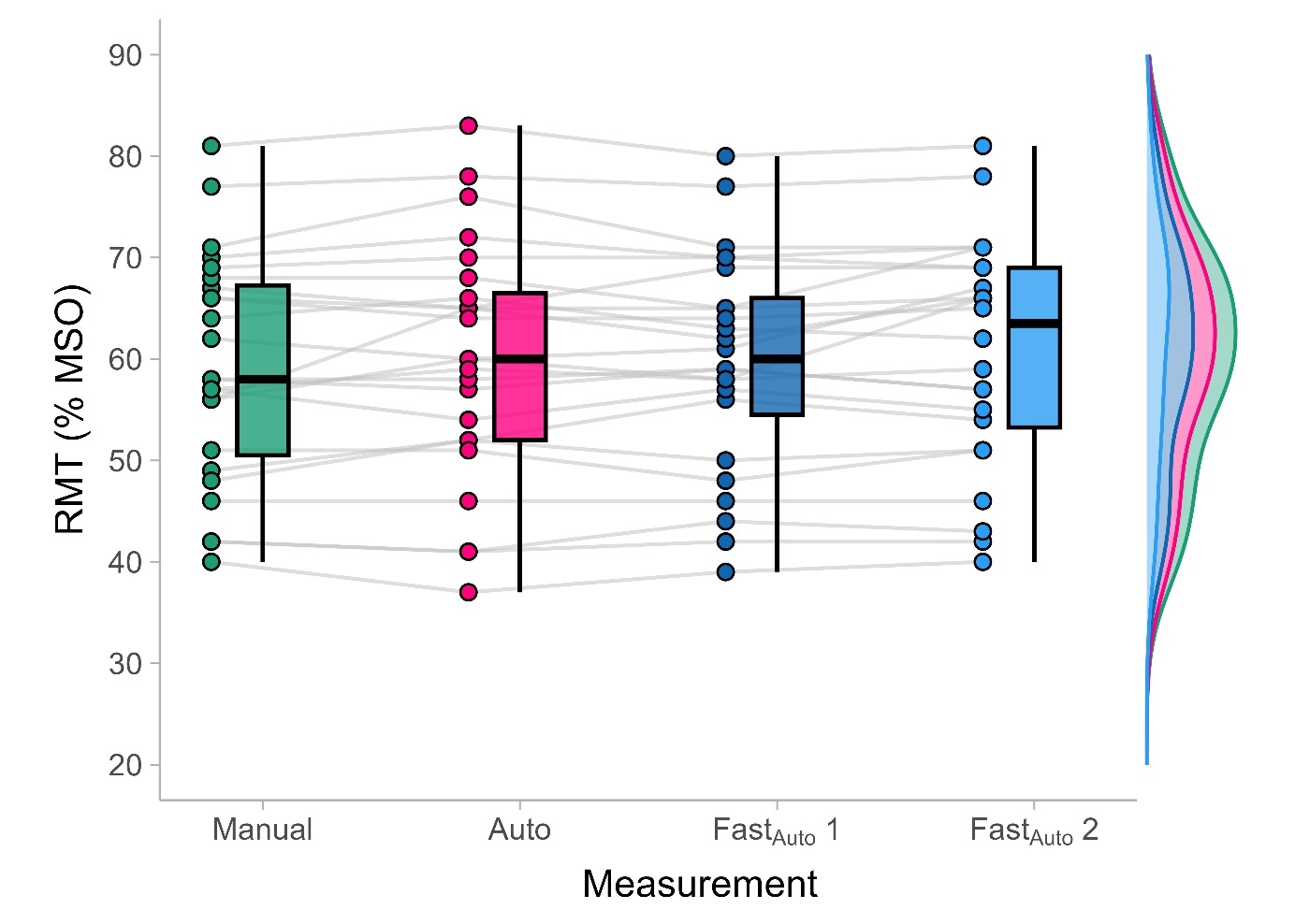

**Supplementary Figure 3.** Boxplots and distributions of the different measurements of the Resting Motor Threshold (RMT), in terms of % of Maximal Stimulator Output (% MSO), in Experiment 2 (N=24).

**Experiment 2: Limits of Agreement**

We again generated Bland-Alman plots for Experiment 2, which are shown in Supplementary Figure 4.

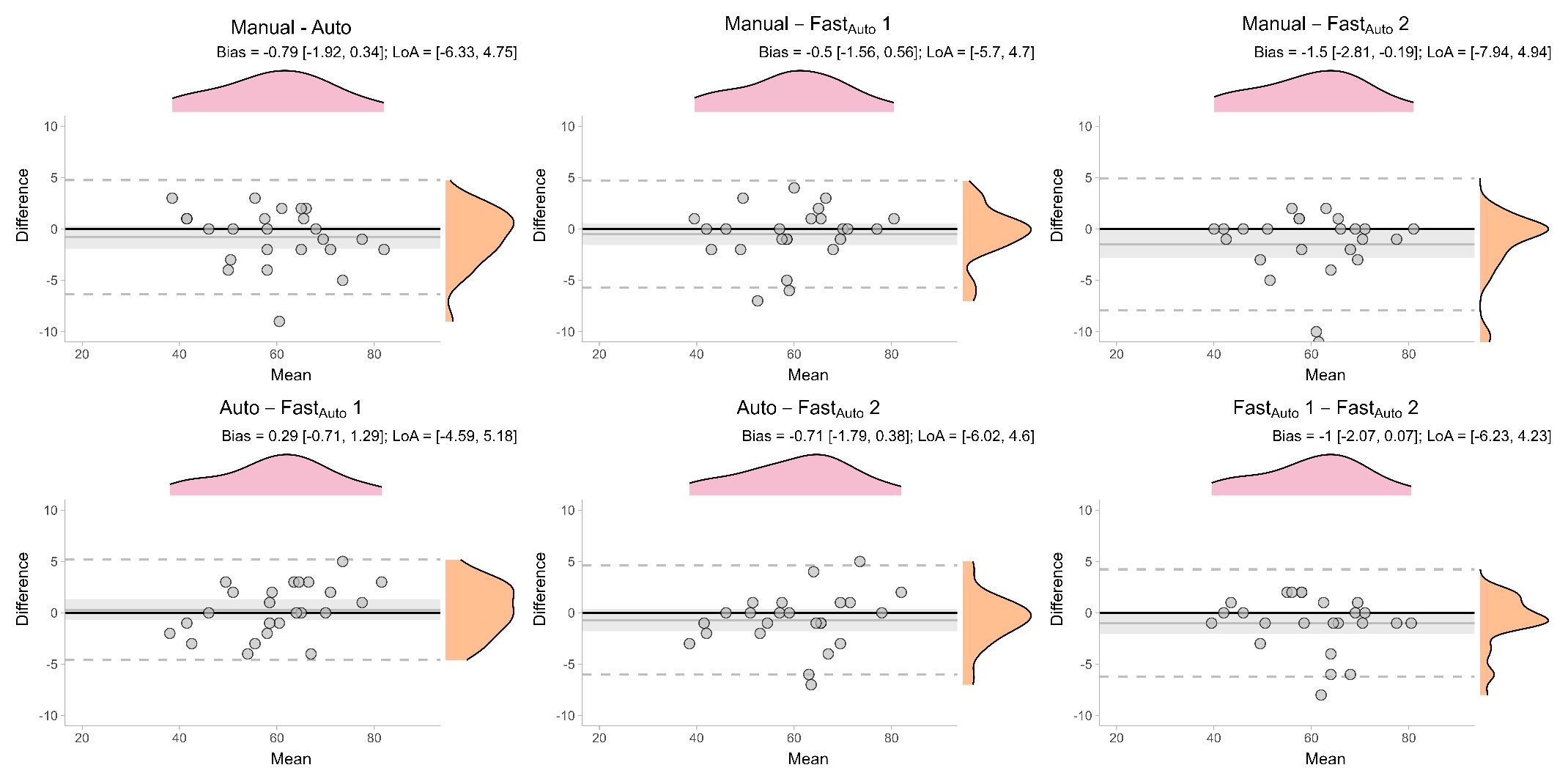

**Supplementary Figure 4.** Bland-Altman plots comparing the manual RMT measurements with both the ‘Auto’ and ‘Fast_Auto_’ versions of the automated procedure for Experiment 2. Each panel displays the mean versus the difference between two measurements. The horizontal black line indicates the mean bias, and shaded areas represent the 95% confidence interval around it. Dashed horizontal lines show the Limits of Agreement (LoA). Bias and Limits of Agreement are reported above each panel.
